# Maternal-fetal genetic interactions, imprinting, and risk of placental abruption

**DOI:** 10.1101/340687

**Authors:** Tsegaselassie Workalemahu, Daniel A. Enquobahrie, Bizu Gelaye, Mahlet G. Tadesse, Sixto E. Sanchez, Fasil Tekola-Ayele, Anjum Hajat, Timothy A. Thornton, Cande V. Ananth, Michelle A. Williams

**Author notes:** Corresponding Author (TW). These authors contributed equally to this work (Tsegaselassie Workalemahu, Daniel A. Enquobahrie). These authors also contributed equally to this work (Mahlet G. Tadesse, Timothy A. Thornton).

## Abstract

Maternal genetic variations, including variations in mitochondrial biogenesis (MB) and oxidative phosphorylation (OP), have been associated with placental abruption (PA). However, the role of maternal-fetal genetic interactions (MFGI) and parent-of-origin (imprinting) effects in PA remain unknown. We investigated MFGI in MB-OP, and imprinting effects in relation to risk of PA. Among Peruvian mother-infant pairs (503 PA cases and 1,052 controls), independent single nucleotide polymorphisms (SNPs), with linkage-disequilibrium coefficient <0.80, were selected to characterize genetic variations in MB-OP (78 SNPs in 24 genes) and imprinted genes (2713 SNPs in 73 genes). For each MB-OP SNP, four multinomial models corresponding to fetal allele effect, maternal allele effect, maternal and fetal allele additive effect, and maternal-fetal allele interaction effect were fit under Hardy-Weinberg equilibrium, random mating, and rare disease assumptions. The Bayesian information criterion (BIC) was used for model selection. For each SNP in imprinted genes, imprinting effect was tested using a likelihood ratio test.

Bonferroni corrections were used to determine statistical significance (p-value<6.4e-4 for MFGI and p-value<1.8e-5 for imprinting). Abruption cases were more likely to experience preeclampsia, have shorter gestational age, and deliver infants with lower birthweight compared with controls. Models with MFGI effects provided improved fit than models with only maternal and fetal genotype main effects for SNP rs12530904 (log-likelihood ratio=18.2; p-value=1.2e-04) in *CAMK2B*, and, SNP rs73136795 (log-likelihood ratio=21.7; p-value=1.9e-04) in *PPARG*, both MB genes. We identified 311 SNPs in 35 maternally-imprinted genes (including *KCNQ1, NPM*, and, *ATP10A)* associated with abruption. Top hits included rs8036892 (p-value=2.3e-15) in *ATP10A*, rs80203467 (p-value=6.7e-15) and rs12589854 (p-value=1.4e-14) in *MEG8*, and rs138281088 in *SLC22A2* (p-value=1.7e-13). We identified novel PA-related maternal-fetal MB gene interactions and imprinting effects that highlight the role of the fetus in PA risk development. Findings can inform mechanistic investigations to understand the pathogenesis of PA.

**Author summary:** Placental Abruption (PA) is a complex multifactorial and heritable disease characterized by premature separation of the placenta from the wall of the uterus. PA is a consequence of complex interplay of maternal and fetal genetics, epigenetics, and metabolic factors. Previous studies have identified common maternal single nucleotide polymorphisms (SNPs) in several mitochondrial biogenesis (MB) and oxidative phosphorylation (OP) genes that are associated with PA risk, although findings were inconsistent. Using the largest assembled mother-infant dyad of PA cases and controls, that includes participants from a previous report, we identified novel PA-related maternal-fetal MB gene interactions and imprinting effects that highlight the role of the fetus in PA risk development. Our findings have the potential for enhancing our understanding of genetic variations in maternal and fetal genome that contribute to PA.

## Introduction

Disturbances that involve mitochondrial biogenesis (MB) and oxidative phosphorylation (OP) underlie pathologic mechanisms leading to placental abruption (PA) - a complex multifactorial disease characterized by premature separation of the placenta from the wall of the uterus [1]. Genome-wide and candidate gene association studies have identified common maternal single nucleotide polymorphisms (SNPs) in several MB and OP genes that are associated with PA risk [2, 3]. However, findings were inconsistent, in line with other previous reports of genetic associations in complex diseases [4, 5].

Investigators have suggested that assessment of maternal-fetal genetic interactions and assessment of imprinting effects, where risk is conferred depending on the parent-of-origin, may explain the missing heritability of complex diseases, particularly those with perinatal origin such as PA [4–6]. PA is a consequence of complex interplay of maternal and fetal genetics, epigenetics, and metabolic factors. For instance, the fetal genome influences placental growth and development, placental implantation and vascularization [7], all of which have been related to PA risk. In addition, many known imprinted genes affect embryonic or trophoblast growth [8] and have been implicated in preeclampsia [9], a known risk factor of PA [10]. Interactions between maternal and fetal genetic variations have previously been demonstrated in preterm delivery – another pregnancy complication with complex origin [11]. However, only one prior study, a study from our group, examined maternal-fetal genetic interactions in relation to risk of PA [2]. Using maternal-placental pairs from 222 PA cases and 198 controls, Denis *et al* reported maternal-fetal genetic interactions on PA risk for two SNPs in the *PPARG* gene (chr3:12313450 and chr3:12412978) and imprinting effects for multiple SNPs in imprinted *(C19MC* and *IGF2/H19)* regions [2]. Using the largest assembled mother-infant dyad of PA cases and controls (503 PA case and 1,052 control mother-infant pairs) to date, that includes participants from the previous report [2], and an expanded set of SNPs (>3,035) in MB-OP genes and imprinted genes, we investigated maternal-fetal genetic interactions and imprinting effects in relation to PA risk.

## Results

Overall PAPE and PAGE study participants (cases and controls) were similar in sociodemographic characteristics and medical/obstetric history (**Table 1**). PAPE and PAGE PA-case mother-infant pairs were similar to control mother-infant pairs with respect to maternal age, marital status, employment, planned pregnancy, infant sex, alcohol use, drug use and vitamin use. Compared to control mother-infant pairs, PA case mother-infant pairs were more likely to smoke during pregnancy, have lower educational attainment, have lower pre-pregnancy body mass index, deliver earlier (i.e., shorter gestational age), deliver infants with lower birth weight, and have a diagnosis of preeclampsia in the current pregnancy.

**Table 1.**
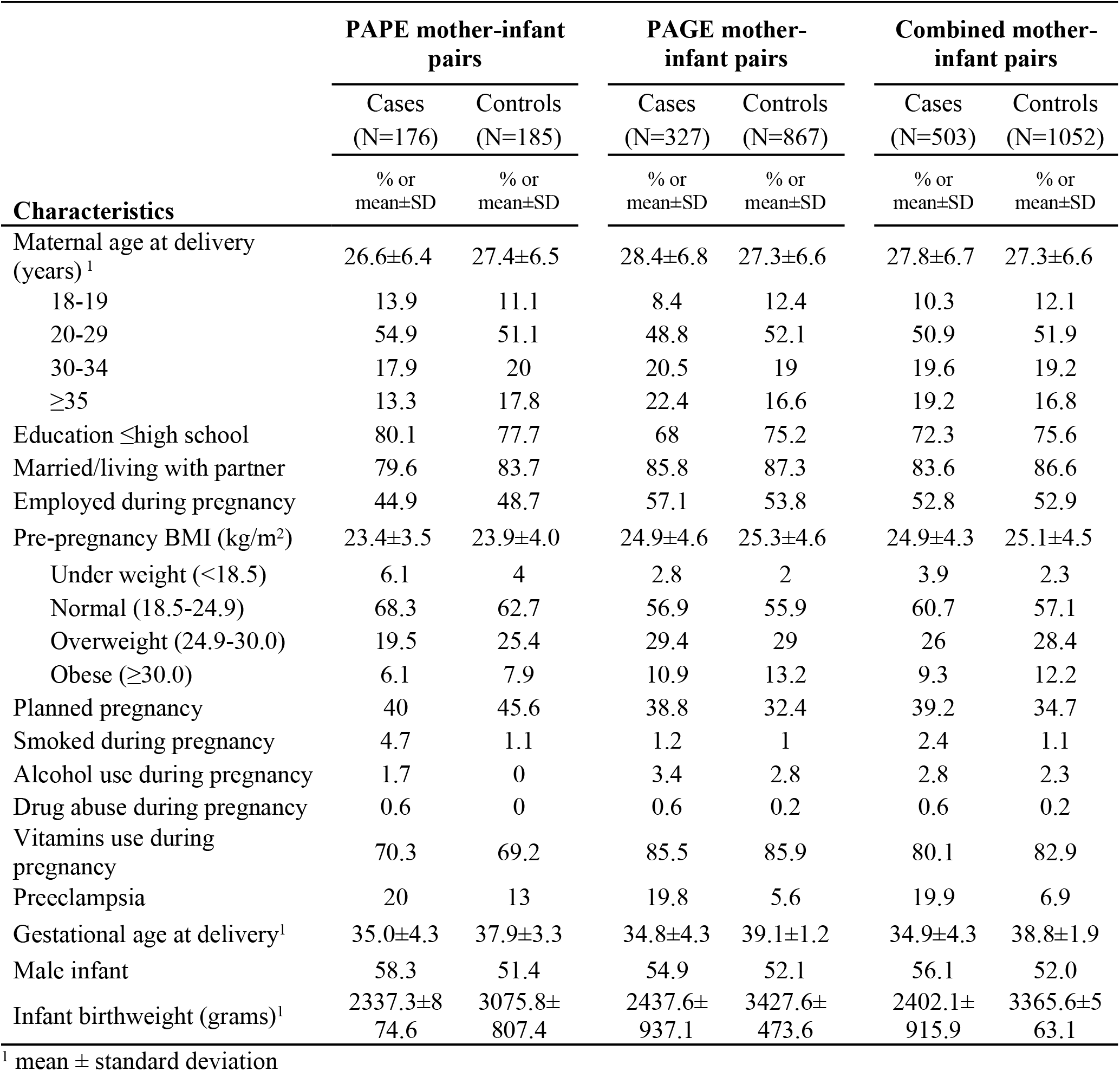
Selected characteristics of study participants

We identified evidence for maternal-fetal genetic interactions on PA risk for two SNPs in two MB genes (**Tables 2** and **3**) where the BIC for Model I was the smallest (**Table 2**): rs12530904 in *CAMK2B* (log-likelihood=-1874.6; p-value=1.2e-04) and rs73136795 in *PPARG* (log-likelihood=-1644.5; p-value=1.9e-04). The risk of PA associated with maternal-fetal genotype combination of GG/AG at the rs12530904 locus was 1.79-fold higher (95%CI: 1.19, 2.69) relative to the maternal-fetal genotype combination of GG/GG. Similarly, the risk of PA associated with maternal-fetal genotype combination of GG/AG at the rs73136795 locus was 2.58-fold higher (95%CI: 1.64, 4.07) relative to the maternal-fetal genotype combination of GG/GG. For two other SNPs, Model I was the best fitting model (**Tables 2** and **3**), although the p-values were not statistically significant after Bonferroni correction: rs12535537 in *CAMK2B* (log-likelihood ratio=12.8; p-value=1.6e-3) and rs35812816 in *PPARG* (log-likelihood ratio=9.0; p-value=0.01). The risk of PA associated with maternal-fetal genotype combination of GG/AG at rs12535537 locus was 1.87-fold higher (95%CI: 1.26, 2.77) relative to GG/GG maternal-fetal genotype combination of rs12535537. The risk of PA associated with maternal-fetal genotype combination of GG/CG at rs35812816 was 1.69-fold higher (95%CI: 1.15, 2.50) relative to the GG/GG maternal-fetal genotype combination.

**Table 2.** SNPs selected with maternal-fetal interaction as best fitting model.

| Gene | rsID | Log Likelihood |  |  |  |  | Likelihood Ratio Test |  |  |  |  | Bayesian Information Criterion (BIC) |  |  |  |
| --- | --- | --- | --- | --- | --- | --- | --- | --- | --- | --- | --- | --- | --- | --- | --- |
|  |  | Model Null | Model F | Model M | Model M+F | Model I | Model F vs Model Null | Model M vs Model Null | Model M+F vs Model M | Model M+F vs Model F | Model I vs Model M+F | Model F | Model M | Model M+F | Model I |
| CAMK2B | rs12535537 | -<br>1975.1 | -<br>1975.0 | -<br>1972.3 | -<br>1972.3 | -<br>1965.8 | 0.30 | 5.66 | 0.10 | 5.46 | 12.84 | 3964.7 | 3959.3 | 3959.2 | 3946.4 |
| CAMK2B | rs12530904 | -<br>1883.9 | -<br>1883.9 | -<br>1879.4 | -<br>1878.8 | -<br>1874.6 | 6.0E-05 | 8.89 | 1.31 | 10.19 | 18.50 | 3782.4 | 3773.5 | 3772.2 | 3763.9 |
| PPARG | rs73136795 | -<br>1655.3 | -<br>1655.3 | -<br>1653.3 | -<br>1653.1 | -<br>1644.5 | 0.02 | 4.12 | 0.43 | 4.54 | 21.72 | 3325.3 | 3321.2 | 3320.8 | 3303.6 |
| PPARG | rs35812816 | -<br>1789.5 | -<br>1766.1 | -<br>1788.5 | -<br>1756.7 | -<br>1752.2 | 46.86 | 2.02 | 63.56 | 18.72 | 8.96 | 3546.8 | 3591.7 | 3528.1 | 3519.1 |

Model F : Fetal only; Model M: Maternal only; Model M+F: Maternal and Fetal only; Model I: Maternal Fetal interaction including the main effects
Likelihood ratio test: 2 x (Log Likelihood of the 1<sup>st</sup> model – Log likelihood of the 2<sup>nd</sup> model)
BIC = -2 x Log Likelihood + #of estimated parameters x ln (#of observations)

**Table 3.** Association estimates of SNPs selected with maternal-fetal interaction as best fitting model.

| Gene | chr:pos | rsID | Risk Allele | Other Allele | Risk Allele Frequency | Fetal Effect Estimates | Maternal Effect Estimates | Interaction Effect Estimates |  |  |
| --- | --- | --- | --- | --- | --- | --- | --- | --- | --- | --- |
| | | | | | | R1 (95% CI) | S1 (95% CI) | $\gamma_{01}$ (95% CI) | $\gamma_{21}$ (95% CI) | P-value |
| CAMK2B | 7:44376785 | rs12535537 | A | G | 0.16 | 1.05<br>(0.87, 1.27) | 1.25<br>(1.04, 1.51) | 1.87<br>(1.26, 2.77) | 1.47<br>(0.85, 2.55) | 1.6E-03 |
| CAMK2B | 7:44377171 | rs12530904 | A | G | 0.14 | 1.00<br>(0.82, 1.22) | 1.34<br>(1.11, 1.61) | 1.79<br>(1.19, 2.69) | 1.18<br>(0.66, 2.12) | 1.2E-04* |
| PPARG | 3:12468410 | rs73136795 | A | G | 0.12 | 0.99<br>(0.79, 1.23) | 0.79<br>(0.63, 0.99) | 2.58<br>(1.64, 4.07) | 0.77<br>(0.27, 2.24) | 1.9E-04* |
| PPARG | 3:12439348 | rs35812816 | C | G | 0.17 | 0.51<br>(0.42, 0.62) | 1.16<br>(0.94, 1.45) | 1.69<br>(1.15, 2.50) | 1.47<br>(0.42, 5.00) | 1.1E-02 |

chr:pos : dbSNP build 37 hg19 chromosome:position
R1: Risk of PA associated with a copy of fetal risk allele (Fetal genotype effect)
S1: Risk of PA associated with a copy of maternal risk allele (Maternal genotype effect)
$\gamma_{01}$ : Risk of PA associated with 1 copy of risk allele from fetus and zero copy of the risk allele from the mother $\gamma_{21}$ : Risk of PA associated with 1 copy of risk allele from fetus and 2 copies of the risk allele from the mother \* statistically significant after Bonferroni correction for multiple testing. p-values correspond to the log-likelihood ratio test comparing the main effects to the interaction effect

We identified 310 SNPs in 31 imprinted genes with imprinting effects (224 maternally expressed, 79 paternally expressed, 3 isoform-dependent, and 4 random) on PA risk that reached statistical significance after Bonferroni correction (**Table 4 and Supplementary Table 1**). These imprinted genes included *KCNQ1* (103 SNPs), *NTM*(30 SNPs), and, *ATP10A* (24 SNPs). Top hits in these analyses were rs8036892 (p-value=2.3e-15) in *ATP10A*, rs80203467 (p-value=6.7e-15) and rs12589854 (p-value=1.4e-14) in *MEG8*, and rs138281088 in *SLC22A2* (p-value=1.7e-13) (**Table 4**).

**Table 4.**
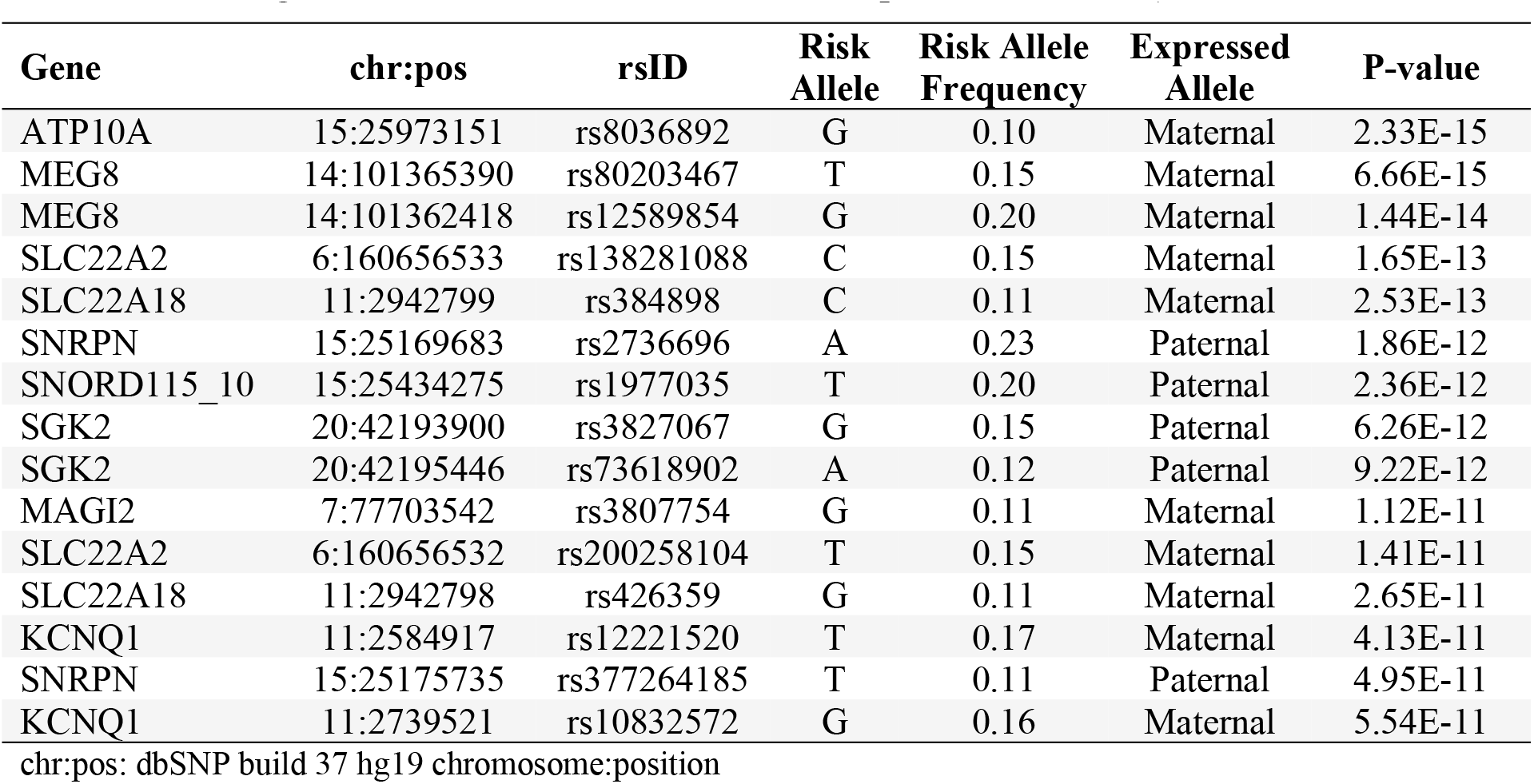
SNPs of imprinted genes with parent-of-origin effect on PA risk (top 15 SNPs out of 311 that were significant after Bonferroni correction [p-value<1.84e-5])

## Discussion

In the current study, we identified several novel maternal-fetal MB SNP interactions and imprinting effects on PA risk. Maternal-fetal interactions were observed for SNPs in *CAMK2B* (rs12530904) and *PPARG* (rs73136795) while potential maternal-fetal interactions were observed for two other SNPs in the same genes (rs12535537and rs35812816 in *CAMK2B* and *PPARG*, respectively). Imprinting effects were observed for 310 SNPs in imprinted genes including *KCNQ1, NTM, ATP10A, MEG8*, and *SLC22A2.*

In the only other similar published study related to PA, our team reported maternal-fetal interaction for two *PPARG* SNPs (chr3:12313450 and chr3:12412978) and imprinting effect for eight SNPs, six in the *C19MC* region and two in *IGF2-H19* [2]. While we found maternal and infant interactions on PA risk for *PPARG* SNP rs73136795 (chr3:12468410), the two previously reported *PPARG* SNPs were not in the set of SNPs we evaluated in the current study as they failed imputation quality (Info<0.3) in the PAGE study. Similarly, we identified imprinting effect on PA risk for *IGF2* SNP rs11564732 (chr1:2150895, p-value=9.3e-06); however, the previously reported SNPs in *C19MC* or *IGF2-H19* genes were not evaluated in the current study because they were not genotyped or imputed in the PAGE study. Other studies have previously investigated interactions between maternal and fetal genetic variations on maternal and infant outcomes [11, 25] [26]. For instance, interaction between maternal and fetal genetic variations at the G308A locus of *TNF-alpha* gene on risk of preterm delivery (PTD) has been reported by a study conducted among 250 PTD and 247 control Han Chinese families [11]. The combined maternal-fetal genotype GA/GA at the locus was associated with reduced risk of PTD (risk ratio=0.20 [95%CI: 0.07, 0.58])[11]. Goddard et. al. [26] observed evidence for maternal-fetal interaction at the rs5742620 loci in *IGF1* gene on preeclampsia risk. Maternal-fetal pairs with

AC/AC genotype at this locus had a 2.4-fold (p-value=0.0035) increased risk of preeclampsia compared to maternal-fetal pairs with CC/CC genotypes [26]. None of the SNPs investigated by the previous investigators were evaluated in our present study. However, collectively, findings from across available studies support possibility of maternal-fetal genetic interactions in pregnancy complications.

Our observation of *PPARG* maternal-fetal interaction on PA risk is noteworthy, not only because we found similar, although in different SNPs, interactions in our previous study, but also because the gene has been well-described in relation to placental growth, development, and function [2, 27, 28]. *PPARG* (peroxisome proliferator-activated receptor gamma) belongs to the PPAR-family of genes and is a master regulator of MB [29]. *PPARG* is highly expressed in the placenta, inhibits trophoblast invasion through oxidized LDL in cytotrophoblasts of cells involved in invasion of the uterus [27, 28], and is associated with the development of preeclampsia [28], an established risk factor of PA [10]. The other gene where we found significant maternal-fetal interactions was *CAMK2B* (calcium/calmodulin-dependent protein kinase [CaM kinase] II beta), a CaMK family gene implicated in contraction-induced regulation of calcium handling in skeletal muscle and MB [30, 31]. *CAMK2B* is among several genes in myometrial relaxation and contraction pathways that are either transcribed in myometrial muscle cells or act upon the myometrium to regulate contraction [32], through pathways that involve oxytocin receptor activation [33]. Preterm uterine contractions are associated with PA risk [34].

We found strong evidence for imprinting effect of several imprinted genes (including *KCNQ1, NTM*, and *ATP10A)* on PA risk. Imprinted genes may affect maternal-fetal interactions that affect placental development [9, 35–37]. Imprinted maternal alleles are required for the development of the embryo, and imprinted paternal alleles regulate formation of the placenta and the surrounding membranes of the embryo [8, 38]. *KCNQ1* encodes a voltage-gated potassium channel required for the repolarization phase of action potential [39]. *KCNQ1* is expressed in the placenta and implicated in embryonic and placental growth [36]. In mice, maternally inherited target deletion of *ASCL2*, a gene that resides in *KCNQ1* cluster, is lethal due to failure of placental formation [37, 40]. The potential roles of both *NTM* and *ATP10A* in pregnancy-related outcomes have not been described before. However, both *NTM* (neurotrimin), involved in neuronal cell adhesion, and *ATP10A*, ATPase phospholipid transporting 10A gene, are expressed in the placenta [41, 42]. The imprinting/parent-of-origin effects of *KCNQ1, NTM*, or *ATP10A* on pregnancy-related outcomes have not been reported before. We found imprinting effect for rs11564732 in *H19*, a maternally imprinted gene near *IGF2*, for which we previously reported similar imprinting effects. *H19/IGF2* regulates the development of the embryo and differentiation of cytotrophoblast cells [43], and was implicated in preeclampsia [44, 45], a known risk factor of PA [10].

In post-hoc exploratory analyses, we examined functions and functional relationships of the 35 imprinted genes that were represented by SNPs with significant imprinting effects using Ingenuity Pathway Analysis (IPA, Ingenuity, Redwood, CA) [46]. In the IPA based on the Ingenuity Pathways Knowledge Base (IPKB), gene-enrichment of networks was assessed using network score, negative log of p-values of a modified Fisher’s exact test. Based on these analyses, the top two enriched gene networks were a network (Score=29) of cell cycle, cell morphology (**Supplementary Table 2** and **Supplementary Fig 1**), and a network (Score=29) of cardiovascular disease and free radical scavenging (**Supplementary Table 2** and **Supplementary Fig 2**). Both of these networks align well with what is known about PA pathogenesis [3].

The underlying genetic architecture of PA has been examined in previous genome-wide [2, 3, 12] and candidate gene association studies [2, 3, 12, 13, 47], which reported predominantly common, non-coding variants with modest effects and limited replication. Important strategies to address subsequent missing heritability include family studies that assess gene-gene interactions and imprinting effects [4, 5]. Family studies are advantageous because affected relatives are more likely to share two nearby epistatic loci in LD that would be unlinked in unrelated individuals [5] and ignoring imprinting effects can mask true associations and diminish the proportion of heritability explained [5]. Other sources of missing heritability may be low frequency variants with intermediate effect. These should be tractable through larger sized studies and imputation of genome-wide data [5].

Our study is the most comprehensive investigation, to date, of maternal-fetal genetic interactions and imprinting effects on PA risk. These findings have the potential for enhancing our understanding of genetic variations in maternal and fetal genome that contribute to PA, a multi-factorial heritable disorder. The study addresses the potential limitations of sample size in previous studies. By conducting 1000 genomes genotype imputations, we analyzed a comprehensive set of SNPs in MB-OP and imprinted genes. However, in the current study, SNPs that did not overlap between PAPE and PAGE were excluded, not allowing us to examine some previously reported SNPs. Other limitations of our study include potential misclassification of sub-clinical PA (i.e. those with less placental disruption and consequent bleeding), which may limit the interpretation of the study results or reduce statistical power. We also did not distinguish between severe and mild cases of PA, which may have different risk factors or underlying mechanisms [48]. Our study was restricted to live births and interaction or imprinting effects that contribute to stillbirth, a common complication of PA, may have been missed. We assessed maternal-fetal genetic interaction on PA risk using MB-OP candidate SNPs. There could be similar interactions in other metabolic functions. This study may still be underpowered for small effects and rare genotypes. Finally, findings from the current study population may not be generalizable to other populations with different population genetic structure or PA risk pattern. Therefore, replication studies in different populations are critical to fully understand maternal-fetal genetic interactions and imprinting effects on PA risk.

In sum, findings in this study confirm the role of interactions between maternal and fetal genetic variations in MB and imprinting effects in PA. These findings highlight the potential of understanding the complex interplay between maternal and fetal genetic factors in explaining the missing heritability of PA and PA-related risk stratification. This may inform potential preventative and therapeutic targets of PA.

## Materials and methods

### Study setting and study populations

The study was conducted among participants of the Peruvian Abruptio Placentae Epidemiology (PAPE) and Placental Abruption Genetic Epidemiology (PAGE) studies, case-control studies of PA conducted in Lima, Peru. Both PAPE and PAGE studies had similar study objectives and study designs, which have been reported before [2, 3, 12, 13]. Briefly, participants were recruited among women who were admitted for obstetrical services to antepartum wards, emergency room, and labor and delivery wards of participating hospitals.

Participants who were less than 18 years of age, delivered infants from a multifetal pregnancy, had medical records that were insufficient to determine the presence or absence of PA (described below), had other diagnoses associated with third trimester bleeding (e.g. placenta previa), or reported taking blood thinning medications were excluded from the studies. PAPE participants who provided maternal blood and placental samples at delivery were included in the current analyses. PAGE participants who provided maternal saliva and newborn buccal cells at delivery were included in the current analyses. The total number of participants included in the current analyses, after exclusions and sample quality control steps (described below), were 176 PA case and 185 control maternal-infant pairs from the PAPE study and 327 PA case and 867 control mother-infant pairs from the PAGE study (a total of 503 PA case and 1,052 control mother-infant pairs). Study protocols of both studies were approved by the Institutional Review Boards (IRBs) of participating institutions and the Swedish Medical Center, Seattle, WA, where the studies were based. All participants provided written informed consent.

### Data collection

PAPE and PAGE study participants were interviewed by trained personnel using standardized, structured questionnaires to collect information on sociodemographic characteristics and risk factors including maternal age, medical history, and smoking (both current and pre-pregnancy). Information on the course and outcomes of the pregnancy were abstracted from maternal medical records. Emergency room admission logbooks, labor and delivery admission logbooks, and the surgery report book were reviewed to determine a diagnosis of PA based on evidence of retro-placental bleeding (fresh blood) entrapped between the decidua and the placenta or blood clots behind the placental margin, accompanied by any two of the following: (i) vaginal bleeding at ≥20 weeks in gestation that is not due to placenta previa or cervical lesions; (ii) uterine tenderness and/or abdominal pain (without other causes, such as those due to hyperstimulation from pitocin augmentation); and, (iii) non-reassuring fetal status or fetal death. Control participants, who did not have a diagnosis of PA in the current pregnancy, were randomly selected from eligible pregnant women who delivered at the participating hospitals during the respective study periods.

In PAPE, maternal blood was obtained for maternal DNA extraction. In addition, placentas were collected immediately after delivery, weighed, double bagged and transported in coolers. Tissue biopsies (approximately 0.5 cm^3^ each) were obtained from 8 sites (4 maternal and 4 fetal) by stripping the chorionic plate and overlying membranes. The biopsy samples were taken from the fetal side and sampled for fetal genomic DNA extraction by placing them in cryotubes, snap frozen in liquid nitrogen, and stored at -80°C until analyses. In PAGE, maternal saliva and newborn buccal cells were collected, plated and stored using the Oragene™saliva cell collection kits (OGR500 and OGR250, DNA Genotek Ottawa, Canada), for DNA extraction and genotyping.

Genomic DNA extraction in the PAPE study were conducted using Gentra PureGene Cell kit (Qiagen, Hilden, Germany). SNP genotyping to characterize genome-wide variations were performed using Illumina Cardio-Metabochip (Illumina Inc., San Diego, CA) platform. In the PAGE study, genomic DNA were extracted using the Qiagen DNAeasy™ system and manufacturer protocols (Qiagen, Valencia, CA). SNP genotyping to characterize genome-wide variations were performed using the Illumina HumanCore-24 BeadChip (Illumina Inc., San Diego, CA) platform.

Maternal and fetal SNP data quality control procedures were applied using identical criteria (in PAPE and PAGE studies) before data analyses. SNPs were excluded if they had excessive missing genotype (SNPs with genotype call rate of <95%), deviated from Hardy-Weinberg equilibrium (HWE; p-value<1e-05), and had low minor allele frequency (MAF<0.05). The total number of SNPs, directly genotyped, that remained for further analyses in PAPE and PAGE studies were 128,371 and 241,301, respectively. Maternal-fetal pairs (PAPE n=23; PAGE n=10) were excluded if they were duplicates or related (Identity by Decent [IBD] value>0.9), had more than 5% of genotyping failure rate (PAPE n=45; PAGE n=51), and had excess heterozygosity rates (outside the range of mean ± 3 standard deviations of heterozygosity rate; PAPE n=5; PAGE n=15). PAGE and PAPE genotype data were then imputed to infer unobserved genotypes using identical steps. The genotype data were phased using SHAPEIT [14] to infer haplotypes and improve imputation accuracy using the 1000 Genomes haplotypes as the reference. Phased haplotypes were then used to impute the non-typed SNPs using IMPUTE2 [15]. After imputation and further quality control (filtering SNPs with imputation certainty score (Info)<0.3, HWE <0.00001, genotyping call rate<0.05, and MAF<0.05), a total of 5,553,176 and 5,314,631 SNPs were available for selection of genes and SNPs (described below) in the PAPE and PAGE studies, respectively.

Candidate genes with described functions in MB and OP were selected from previously published studies [3, 16–21]. Among 785 (in 101 MB-OP genes) and 359 SNPs (in 26 MB-OP genes) that were genotyped/imputed in the PAPE and PAGE studies, respectively, 322 overlapping SNPs (in 24 MB-OP genes) were selected. Pair-wise linkage disequilibrium (LD) was assessed between these 322 SNPs using SNAP [22]. A total of 78 independent SNPs (LD<0.80 in the set) in the 24 MB-OP genes (see Supplementary Table 1) were selected for maternal-fetal interaction analyses. Similarly, a total of 12,459 SNPs in 83 imprinted genes from PAPE study and 10,030 SNPs in 78 imprinted genes from PAGE study were identified using GeneImprint [23]. Out of 9,666 SNPs in 73 imprinted genes that overlap between the two studies, a total of 2,713 independent SNPs (in 35 imprinted genes) were selected for imprinting effect analyses.

### Statistical analyses

Mean and standard deviations for continuous variables and proportions for categorical variables were used to compare the characteristics of PA case and control participants. In maternal-fetal interaction analyses, for each SNP, similar to Denis et. al. [2], four models corresponding to allele effects operating only at fetal level (Model F), allele effects operating at maternal level (Model M), an additive model of maternal and fetal effects (Model M+F), and a model that includes a maternal-fetal interaction effect (Model I) were considered. For the latter, we applied a parametrization that introduces two interaction terms capturing incompatibility between maternal and fetal genotypes; the interaction effects operate when the infant has one copy and the mother has either zero or two copies of the risk allele. The Bayesian information criterion (BIC) was used for model selection. In addition to the maternal and fetal genotype effects, we estimated the risk ratio (RR) of disease when the infant has one copy and the mother has zero copies of the risk allele. The reference group is defined as mother-infant pairs carrying zero copies of the risk allele.

Imprinting/parent-of-origin effect, which corresponds to the factor multiplying the disease risk if the infant inherits a risk allele from the mother, was assessed using a likelihood ratio test [2]. Bonferroni corrections were applied to correct for multiple testing: p-value<6.4e-4 for the 78 maternal-fetal interaction tests, and, p-value<1.8e-5 for 2,713 imprinting effect tests.

All analyses were conducted based on HWE, random mating, and rare disease assumptions. Statistical analyses were conducted using PREMIM [24], EMIM [24], R (version i386 3.1.2) and SAS (Version 13).

## Acknowledgment

The authors are indebted to the participants of the PAPE and PAGE studies for their cooperation. They are also grateful to Ms. Elena Sanchez and the dedicated staff members of Asociacion Civil Proyectos en Salud (PROESA), Peru for their expert technical assistance with this research.

## Supporting Information Legends

**Supplementary Table 1**. Mitochondrial biogenesis and/or oxidative phosphorylation pathway genes evaluated in the current study.

**Supplementary Table 2**. SNPs of imprinted genes with imprinting effect on PA risk (310 that were significant after Bonferroni correction [p-values<1.5e-5]).

**Supplementary Table 3**. Networks represented by 35 imprinted genes identified for imprinting effect on PA risk.

**Supplementary Figure 1**. Significant networks (P-value=1.0e-28) represented by 12 imprinted genes with imprinting/parent-of-origin effect on PA risk. Molecules highlighted in purple represent cell cycle and cell morphology pathway.

**Supplementary Figure 2**. Significant networks (P-value=1.0e-29) represented by 12 imprinted genes with imprinting/parent-of-origin effect on PA risk. Molecules highlighted in purple represent cardiovascular disease, cell morphology, and free radical scavenging pathway.

